# TWISP: A Transgenic Worm for Interrogating Signal Propagation in *C. elegans*

**DOI:** 10.1101/2023.08.03.551820

**Authors:** Anuj Kumar Sharma, Francesco Randi, Sandeep Kumar, Sophie Dvali, Andrew M Leifer

**Affiliations:** Department of Physics, Princeton University, Princeton, NJ, 08544; Princeton Neuroscience Institute, Princeton University, Princeton, NJ, 08544

**Keywords:** functional connectivity, calcium imaging, optogenetics, GUR-3, PRDX-2, Dexamethasone, drug-inducible gene expression, *C. elegans*, neurons

## Abstract

Genetically encoded optical indicators and actuators of neural activity allow for all-optical investigations of signaling in the nervous system. But commonly used indicators, actuators and expression strategies are poorly suited for systematic measurements of signal propagation at brain scale and cellular resolution. Large scale measurements of the brain require indicators and actuators with compatible excitation spectra to avoid optical crosstalk. They must be highly expressed in every neuron but at the same time avoid lethality and permit the animal to reach adulthood. And finally, their expression must be compatible with additional fluorescent labels to locate and identify neurons, such as those in the NeuroPAL cell identification system. We present TWISP, a Transgenic Worm for Interrogating Signal Propagation, that address these needs and enables optical measurements of evoked calcium activity at brain scale and cellular resolution in the nervous system of the nematode *Caenorhabditis elegans*. We express in every neuron a non-conventional optical actuator, the gustatory receptor homolog GUR-3+PRDX-2 under the control of a drug-inducible system QF+hGR, and calcium indicator GCAMP6s, in a background with additional fluorophores of the NeuroPAL cell ID system. We show that this combination, but not others tested, avoids optical-crosstalk, creates strong expression in the adult, and generates stable transgenic lines for systematic measurements of signal propagation in the worm brain.

## INTRODUCTION

A fundamental goal of neuroscience is to understand how neural signals flow through the brain to process information and generate actions. Genetic model systems are crucial for this understanding, in part, because they provide a platform to express genetically encoded optical indicators and actuators for measuring and manipulating neural activity (Simpson and Looger 2018; Randi and Leifer 2020). Genetically encoded calcium or voltage indicators combined with light-gated ion channels have enabled all-optical functional investigations of neural signaling in circuits or sub-brain regions (Guo *et al*. 2009; Rickgauer *et al*. 2014; Emiliani *et al*. 2015; Franconville *et al*. 2018). Now there is interest in performing such investigations at nervous-system scale.

The *Caenorhabditis elegans* (*C. elegans*) nervous system already has a well-mapped anatomical wiring diagram, called a connectome (White *et al*. 1986; Cook *et al*. 2019; Witvliet *et al*. 2021) and a cell-resolved atlas of gene expression (Taylor *et al*. 2021). Adding measurements of how neural signals propagate at brain scale and cellular resolution would allow for systematic comparisons between a nervous system’s structure, gene expression and function. But commonly used combinations of indicators, actuators and expression strategies are poorly suited for measuring neural signal propagation at brain scale and cellular resolution. Many approaches suffer from unwanted neural activation during imaging due to spectral overlap. For example, 488 nm light typically used to image calcium activity with GCaMP (Chen *et al*. 2013) will activate Chrimson at ∼35% of its on-peak photocurrents (Klapoetke *et al*. 2014). And to our knowledge, light-gated ion channel expression has not been previously reported in every single neuron in a brain, possibly because, as we discover here, high expression throughout the brain can be toxic and sometimes lethal.

An approach is needed with the following requirements: An indicator and actuator pair are needed that avoid spectral overlap, so that neural activity can be imaged without unwanted neural activation. Sufficiently high expression of the actuator and indicator is needed to allow robust measurements while still avoiding lethality. And their expression must be compatible with fluorescent reporters to identify each neuron with respect to the connectome (Yemini *et al*. 2021).

In this work we tested various combinations of calcium indicator, neural actuators, and expression strategies in the nematode *C. elegans*. We present TWISP, a **<u>T</u>**ransgenic **<u>W</u>**orm for **<u>I</u>**nterrogating **<u>S</u>**ignal **<u>P</u>**ropagation and demonstrate this strain’s suitability for large-scale signal propagation mapping of the brain.

## RESULTS

### GUR-3+PRDX-2 and TsChR have excitation spectra compatible with GCaMP imaging

We sought a genetically encoded neural activity indicator and optogenetic actuator that could be co-expressed in each cell to provide optical access to measure and manipulate neural activity of all neurons (Fig. 1a). We focused on calcium indicators because recent families of calcium indicators have typically had brighter fluorescence with larger signal-to-noise ratios than commonly used voltage indicators. The GCaMP family of calcium indicators is the most widely used (Chen *et al*. 2013) and has a single-photon absorbance peak at approximately 498 nm (Fig. 1b). But wavelengths used to excite GCaMP near this peak also excite common optogenetic proteins including ChR2 and Chrimson (Fig. 1c). For example, Chrimson is reported to be excited to roughly ∼35% of its maximal photocurrents by 488 nm light commonly used to image GCaMP (Klapoetke *et al*. 2014). This optical crosstalk leads to unwanted optogenetic stimulation and poses a challenge for single photon calcium imaging. 2-photon imaging of GCaMP provides alternative strategies for avoiding unwanted optical activation (Rickgauer *et al*. 2014; Emiliani *et al*. 2015; Packer *et al*. 2015), but we specifically sought a single-photon imaging solution in order to use spinning-disk confocal microscopy because of that method’s speed and relative ease of use (Nguyen *et al*. 2016). Red shifted calcium indicators avoid spectral overlap with opsins such as ChR2 (Dana *et al*. 2016), but in our hands neither jRGECO1a nor jRCaMP1b (Dana *et al*. 2016) were sufficiently bright for the fast volumetric imaging needed here (short 20 ms camera exposures). We therefore explored blue-shifted optogenetic actuators that might work better with GCaMP.

**Figure 1.**
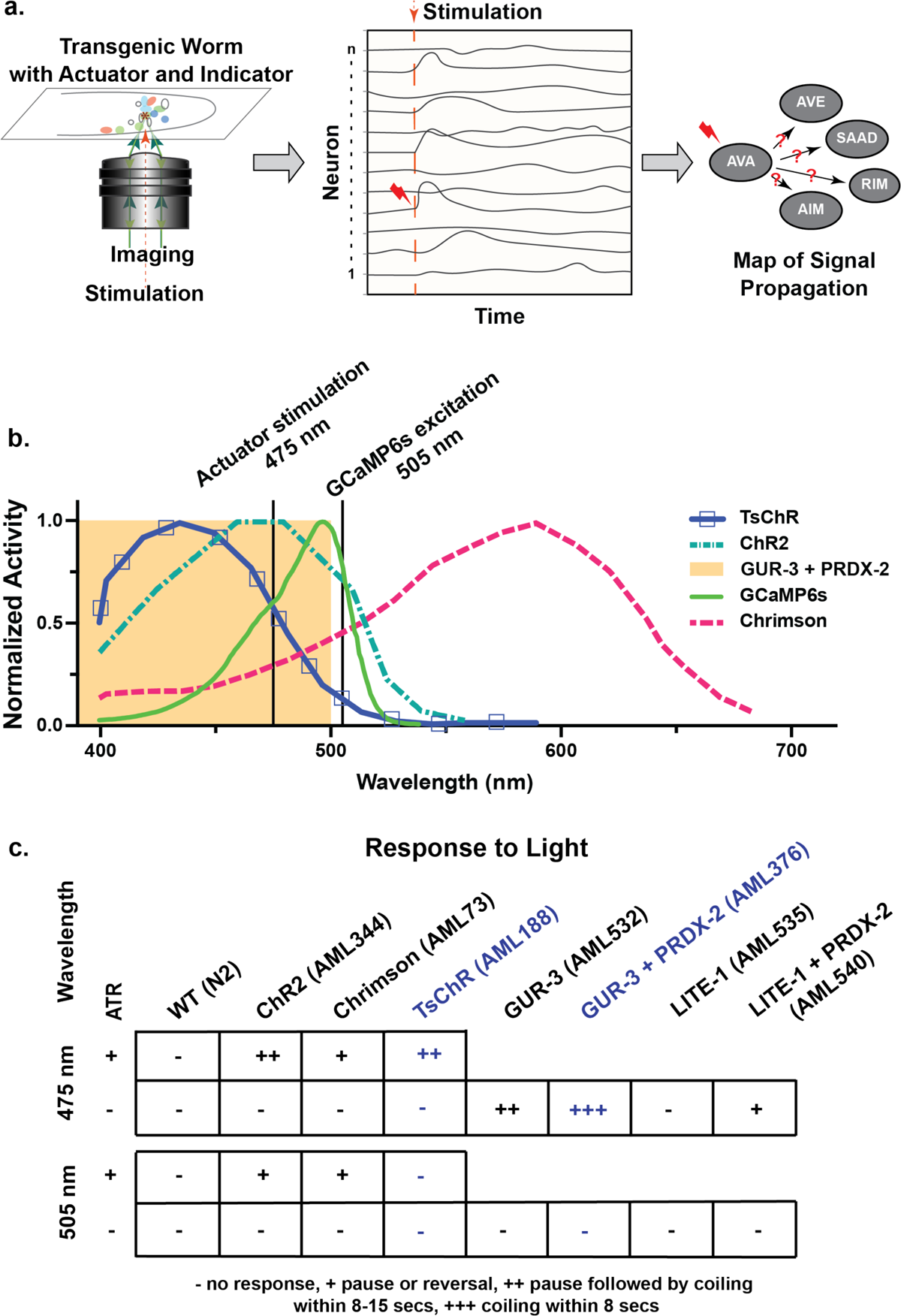
Strategy for all-neuron random-access stimulation and calcium imaging without optical cross talk. a.) Schematic of a transgenic *C. elegans* for measuring signal propagation via optogenetic stimulation and calcium imaging. b.) Previously reported action spectra for several neural actuators, compared with the absorbance spectra of GCaMP6s also shown for comparison. Adapted from: (Chen *et al*. 2013; Husson *et al*. 2013; Klapoetke *et al*. 2014; Bhatla and Horvitz 2015; Dana *et al*. 2016). 505 nm light is close to GCaMP6s absorbance peak at 498 nm. c.) Optogenetic proteins were expressed in every neuron under a *rab-3* promotor. Behavior response to 1.5 mW/mm^2^ illumination of either 505 nm or 475 nm is shown. For animals expressing rhodopsin-based optogenetic proteins, behavior is measured with and without the necessary co-factor all-trans retinal (ATR). Actuators in blue show good compatibility with GCaMP imaging wavelengths.

We tested several actuators (Fig. 1c) and focused on two that were blue-shifted compared to Channelrhodopsin: the opsin TsChR (Klapoetke *et al*. 2014; Farhi *et al*. 2019) and a light-sensitive gustatory receptor homolog GUR-3 that is endogenous to the worm (Bhatla And Horvitz 2015). Both are reported to be activated efficiently by wavelengths near 475 nm light, but less so by 505 nm light (Klapoetke *et al*. 2014; Bhatla And Horvitz 2015). Crucially, 505 nm light is close to GCaMP’s excitation peak of 498 nm and is expected to enable GCaMP imaging with similar efficiency to commonly used 488 nm light (Chen *et al*. 2013). We expressed the optogenetic proteins TsChr or GUR-3 in every neuron under a pan-neuronal *rab-3* promotor. For TsChR we grew the worm on the necessary cofactor all-trans retinal (ATR). For GUR-3, we co-expressed an associated peroxiredoxin, PRDX-2, thought to improve the efficiency of its light response (Fig. 1c). Illuminating animals expressing TsChR or GUR-3+PRDX-2 with 1.5 mW/mm^2^ intensity 475 nm light caused the animals to pause and coil (Fig. 1c). In contrast, illuminating animals with 1.5 mW/mm^2^ intensity 505 nm, near GCaMP’s excitation peak, caused little or no response. In this work we use a hand-scoring method on a four-point scale to characterize the animal’s light responses. In the highest score, “+++,” animals typically coiled within 8 secs of illumination. In the next highest score, “++,” animals typically paused for at least 8 secs and only later coiled. In the second lowest score, “+,” animals paused or reversed but did not coil. And in the lowest score animals rarely responded at all, “-.”, These measurements suggest that both TsChR and GUR-3+PRDX-2 are candidates for co-expression with GCaMP, and that GCaMP can be imaged without unwanted activation of TsChR or GUR-3+PRDX-2.

WT animals did not noticeably pause or coil in response to illumination, as expected, and neither did opsin-expressing animals that lacked the necessary cofactor all-trans retinal (ATR) (Fig. 1c). Also as expected, actuators that rely on peroxiredoxin, such as GUR-3, were most response when PRDX-2 was co-expressed (Liu *et al*. 2010; Bhatla And Horvitz 2015; Gong *et al*. 2017; Quintin *et al*. 2022). Based on these measurements, we concluded that TsChR and GUR-3+PRDX-2 were the two most promising candidates compatible with CGaMP6s for random-access optical activation and imaging, and we proceeded to explore them further. Later in this work we also used a related TsChR variant, eTsChR, that is optimized for more efficient trafficking to the membrane but is otherwise similar (Farhi *et al*. 2019).

### Constitutive overexpression of pan-neuronal TsChR or GUR-3+PRDX-2 is lethal

To be able to manipulate and record activity from every neuron in a single animal, we desire expression of both actuator and indicator in every neuron. The expression level of the actuator must be sufficiently high to evoke a strong response to illumination. We explored different expression levels of the actuators by injecting different concentrations of DNA plasmids for either TsChR or GUR-3+PRDX-2, each with a fluorescent protein (tagBFP or tagRFP, respectively) via co-injectable marker or a SL2 splice site. Higher concentration injections were lethal, but at injection concentrations of 50 ng/µl or below we were able to generate lines that carried the actuator and fluorescent protein in an extrachromosomal array (Fig. 2a). For example, the 35 ng/µl injection concentration GUR-3+PRDX-2 showed strong responses (Fig. 1c). We observed variability in expression levels between animals, as expected for extrachromosomal arrays. For TsChR expressing animals, most animals had very dim expression or no expression of the co-expressed tagBFP reporter and had long generation time, a potential sign of toxicity. Rarely did we observe worms that had high expression. L1s with high expression never developed into adulthood. By comparing observations of the animal’s fluorescent expression with its behavior responses to blue light, we confirmed that higher expression is needed to achieve more robust neural activation. The rare L4s with bright co-expressed BFP responded to 475 nm light by pausing and then coiling (Fig. 2b, Fig. 1c), while animals from the same strain with lower expression, which were more common, merely paused.

**Figure 2.**
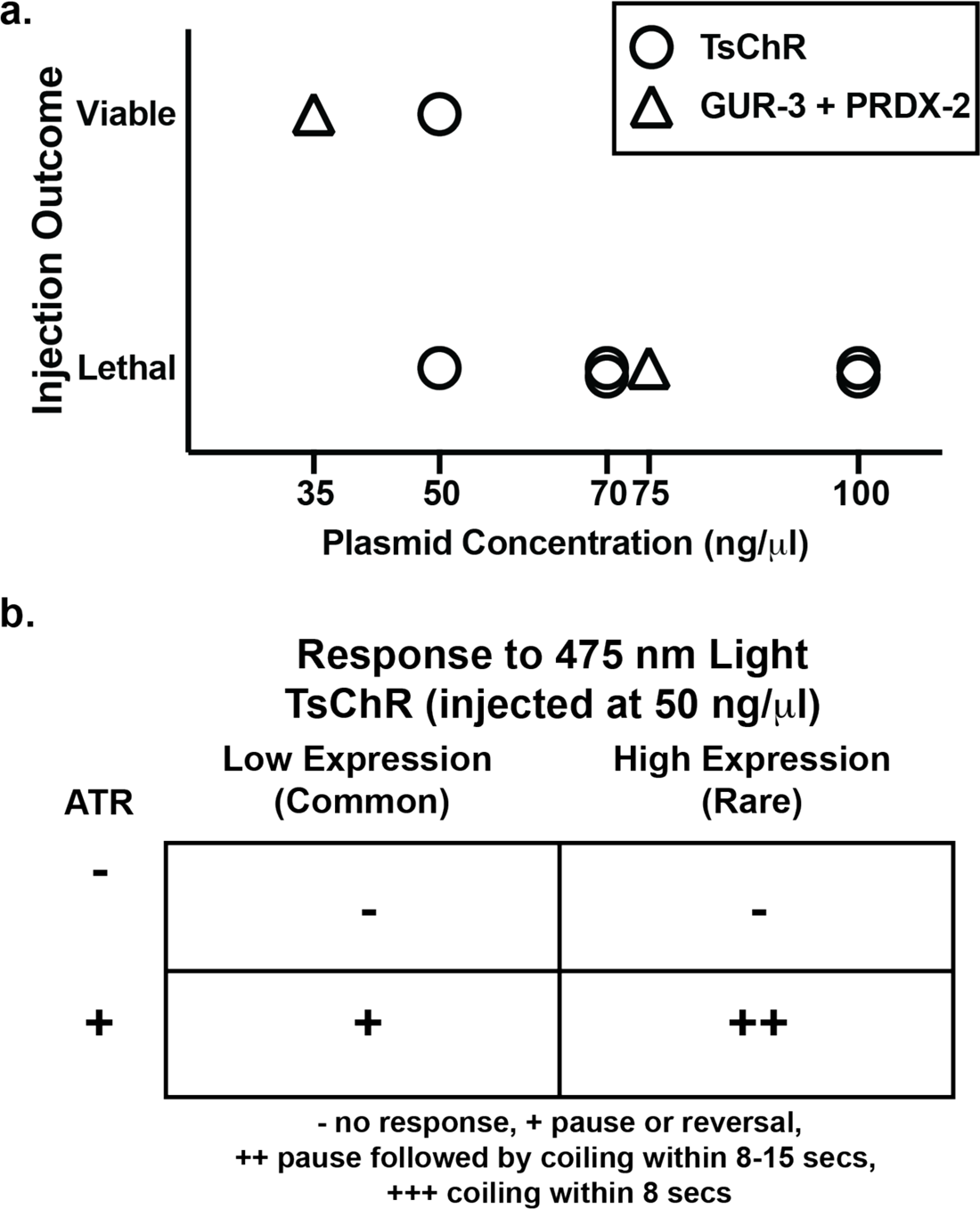
High concentration injections of actuator-containing plasmids are not viable for transgenesis, but higher expression is desirable. a.) Injection concentration for plasmids containing either GUR-3+PRDX-2 or TsChR and the viability of transgene expressing progeny is shown. b.) Light-evoked behavior response of worms expressing TsChR (50 ng/µl injection) in all neurons upon 1.5 mW/mm^2^ 475 nm blue light illumination, with and without the cofactor ATR (All-trans-retinal). Most worms had very low expression, based on fluorescent reporter marker. Rare worms with high expression showed stronger responses.

To avoid variability, we sought to create stable transgenic lines. We attempted to generate integrated strains that would carry the TsChR array in the genome (50 ng/µl), but both classical UV (Evans 2006) and miniSOG-assisted blue-light integration methods (Noma And Jin 2018) failed to create stable lines. Our earlier observation that L1-stage extrachromosomal animals never developed to adulthood suggests that there may be toxicity or arrested development from strong expression of the actuator. The challenges we observed in attaining an integrated strain, and the observations that bright expressing L1s fail to reach adulthood, together suggest that sufficiently high expression of pan-neuronal TsChR or GUR-3+PRDX-2 is lethal or disrupts development.

We and others use the *rab-3* promoter to drive pan-neuronal expression of transgenes (Tursun *et al*. 2011; Nguyen *et al*. 2016; Venkatachalam *et al*. 2016; Yemini *et al*. 2021). *rab-3* mRNA expression peaks early in development: expression is highest during L1 larval stage and then decreases steadily and is lowest in adulthood (Davis *et al*. 2022). We hypothesized that the toxicity from overexpressing our actuator may be highest early in development, and that toxicity from this high expression may act as a bottleneck preventing high expressing lines from propagating. This would explain why we observe bright L1 expression in animals that fail to develop, and why we faced challenges in generating stable integrated lines. If this hypothesis were correct, we reasoned that turning off expression early in development may allow for the generation of stable integrated lines with higher expression later in development. We therefore sought to temporally control expression of our optogenetic actuators.

### Temporal control of actuator expression via QF+hGR > QUAS allows for creation of stable lines with inducible high expression

We set out to temporally control pan-neuronal expression of our optogenetic actuator by using the dexamethasone-inducible QF+hGR > QUAS expression system (Monsalve *et al*. 2019). This system introduces a heterologous gene expression system (Q system) under the control of an exogenous human glucocorticoid receptor binding site. Application of the drug dexamethasone activates an engineered protein, QF-hGR, which then binds to the “QUAS” DNA sequence and activates downstream gene expression.

We designed two sets of multiple plasmids to express either eTsChR or GUR-3+PRDX-2 under the control of the dex-inducible system, which in turn is expressed under the pan-neuronal promoter, *rab-3* (Supplementary Table S2). We injected these plasmids at concentrations of 75 to 100 ng/µl (Fig. 3a), higher than what had previously been viable in the experiments shown in (Fig. 2a). Notably, we obtained lines expressing the QF+hGR regulated transgenes, either eTsChR (AML438) or GUR-3+PRDX-2 (AML376), verified by a co-expressed tagBFP, that survived through adulthood. Animals that were transferred onto dexamethasone containing plates (200 µM dexamethasone in NGM media) as L4s and cultivated overnight showed stronger light evoked responses than similarly aged animals cultivated without dexamethasone, as expected (Fig. 3b). The dexamethasone treated animals exhibited responses similar in strength to the highest-expressing animals of the strain that lacked the temporal drug-inducible expression system (compare Fig. 3b to Fig. 2a). Importantly, for the QF+hGR>GUR-3+PRDX-2 strain on dex, the vast majority of animals exhibited strong responses. In contrast, few animals had expressed strong responses from the non drug-inducible strain. We therefore concluded that temporal control of our optogenetic actuator allows for more consistently strong expression of our actuator in adulthood.

**Figure 3.**
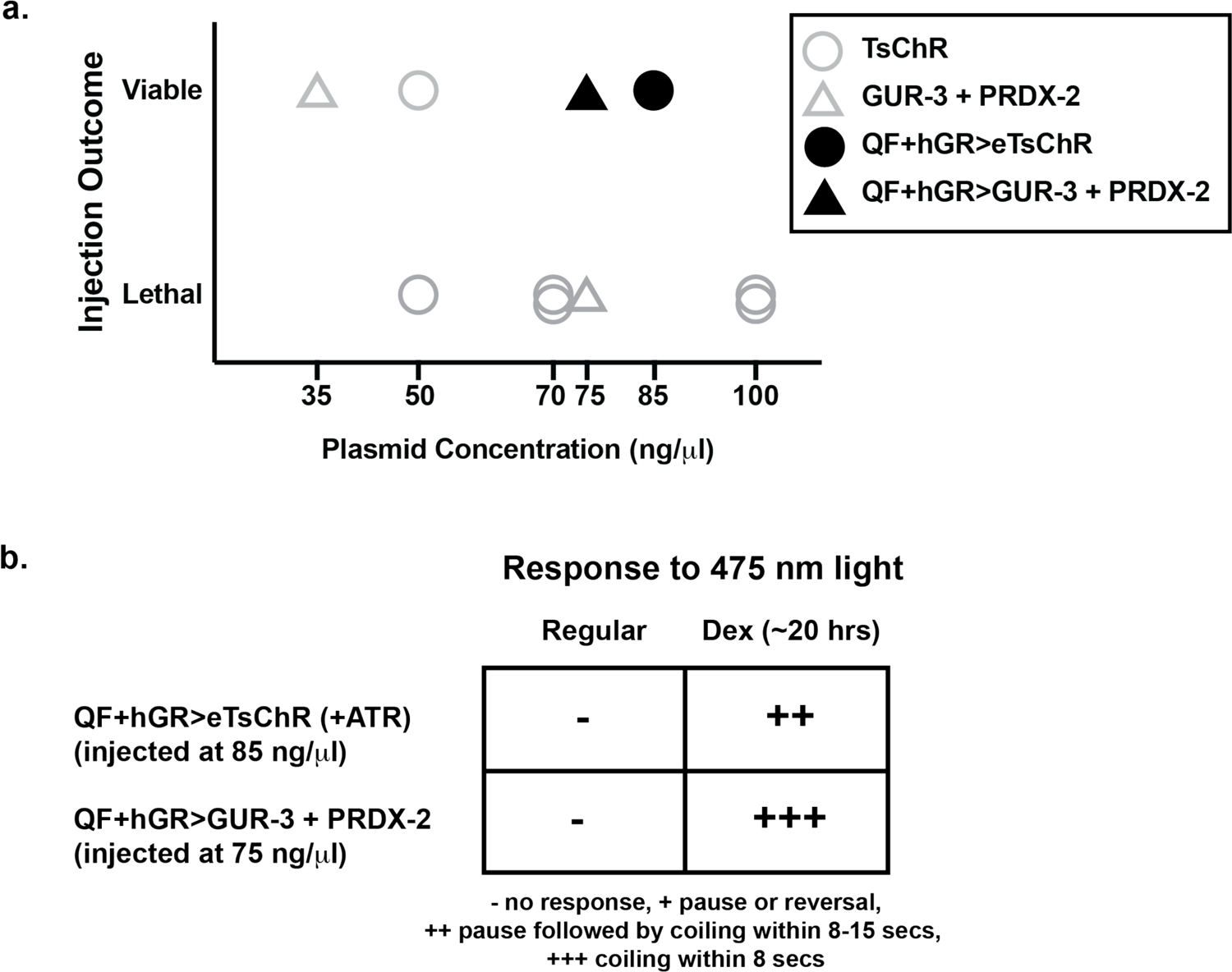
Drug inducible expression enables robust light response while avoiding lethality. a.) Injections of plasmids containing actuators under the control of the QF+hGR drug inducible expression system are viable at higher injection concentrations (black filled shapes) than injections of plasmids for direct expression of the actuators (gray open shapes, same as Fig 2). b.) Exposure to the drug dexamethasone (Dex) evokes actuator expression and confers robust light-response to 475 nm illumination.

The drug-inducible GUR-3+PRDX-2 strain showed slightly stronger light evoked responses than a drug-inducible eTsChR strain (Fig. 3b). The GUR-3+PRDX-2 expressing animals often coiled within 8 secs of blue light illumination, while the eTsChR expressing animals typically only coiled after a longer delay. We therefore generated a stable integrated strain, AML456, via UV integration, that expresses GUR+PRDX-2 under the drug-inducible system, along with a GFP coelomocyte marker, and outcrossed with N2 eight times.

The temporally inducible animals cultivated on dexamethasone for their entire lives showed growth retardation, had notably longer generation times, and visually appeared sick after the second generation (Fig. 4). Drug-inducible animals on chronic dexamethasone qualitatively resembled the non-inducible actuator expressing animals from (Fig. 2). Interestingly progeny of drug-inducible animals cultivated on dexamethasone reverted back to improved health and more normal generation times if they were grown off of dexamethasone (Fig. 4). These observations are consistent with our hypothesis that the pan-neuronal expression of actuator creates a toxicity bottleneck, likely in early adulthood. We conclude that the drug inducible system allows stable transgenic lines to be generated and propagated in a healthy and more timely manner by avoiding this toxicity bottleneck.

**Figure 4.**
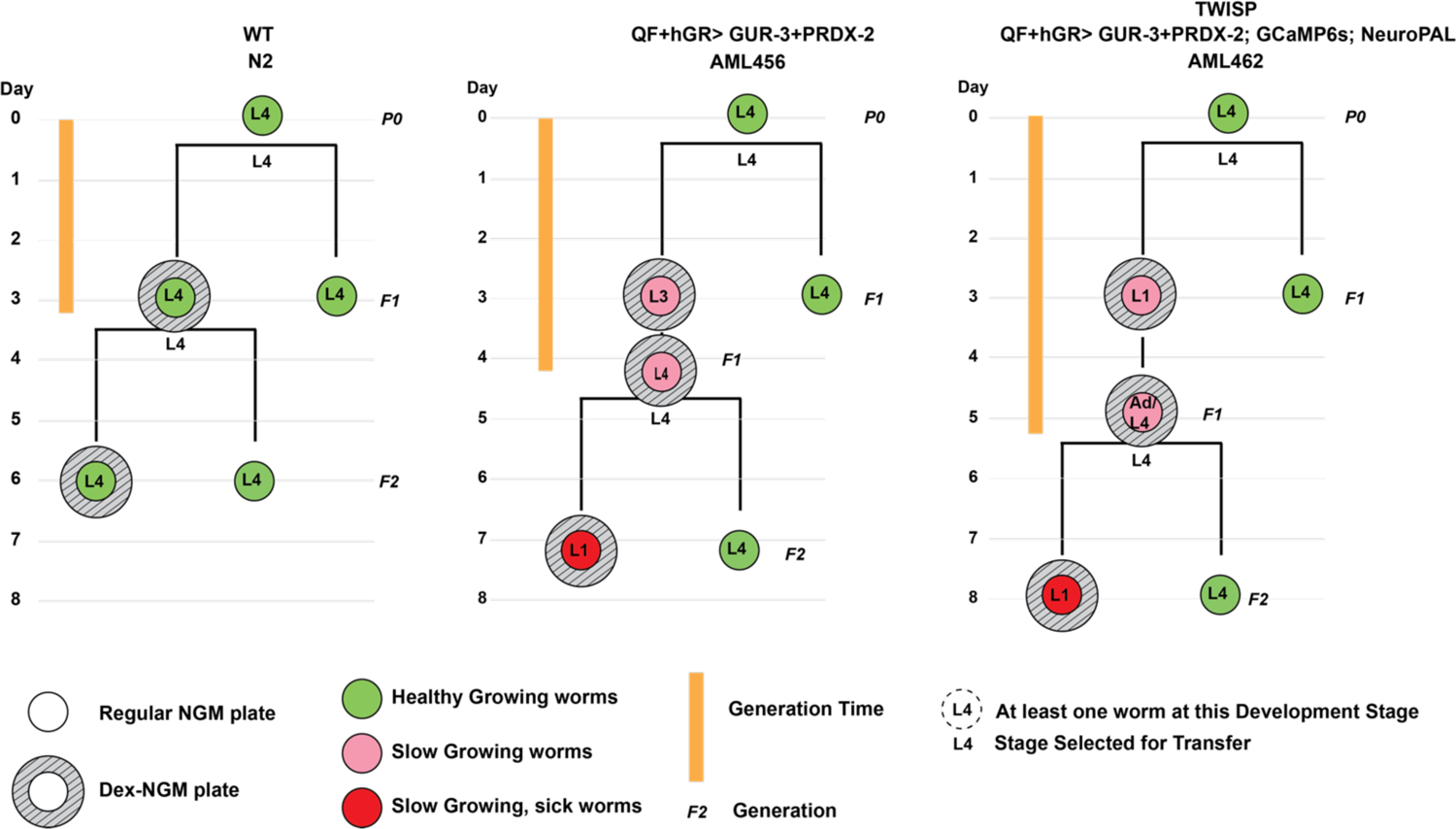
Drug induced actuator expression modulates health and growth. Animals are observed as they are propagated on plates across multiple generations either on or off the drug dexamethasone. Observations of the animal’s health and developmental stage are made at time points indicated by the circles. For each observation, the developmental stage of the most developmentally advance animal found on the plate is reported. Health and generation times improve for the progeny of animals that had previously been exposed to the drug.

### TWISP- a Transgenic Worm for Interrogating Signal Propagation

To generate a strain for measuring signal propagation, we sought an integrated line that combined our drug-inducible actuator system with a calcium indicator and with the NeuroPAL multi-color fluorescent system for neuronal identification (Yemini *et al*. 2021). The NeuroPAL system expresses multiple fluorophores combinatorially in each neuron to enable identification of each neuron with respect to the connectome. The system works by converting each neuron’s gene expression profile into a genetically encoded fluorescent color code. We crossed our drug-inducible actuator line, AML456, into a NeuroPAL+GCaMPs line, AML320 (Yu *et al*. 2021) to create TWISP, a transgenic worm for interrogating signal propagation, AML462. TWISP contains around 50 plasmid constructs integrated into the genome. Each neuron express GCaMP6s; tagRFP; some combination of tagBFP2, cyOFP, and/or mNeptune; and, upon dexamethasone treatment, GUR-3 and PRDX-2. Because NeuroPAL strains had previously been reported to be less active (Yemini *et al*. 2021), we sought to characterize changes in the animal’s vitality and behavior upon the addition of so many genetic components. We therefore measured behavior and lifespan in several transgenic lines, (Fig. 5 and 6). TWISP is noticeably slower, less active, produces animals that have shorter body lengths, and exhibit longer generation time and produce fewer progeny per unit time than wild-type animals. Notably, TWISP is similar in most of these features to our 14x outcrossed NeuroPAL + GCaMP6s strain, AML320 (Fig. 5 and 6) (Yu *et al*. 2021).

**Figure 5.**
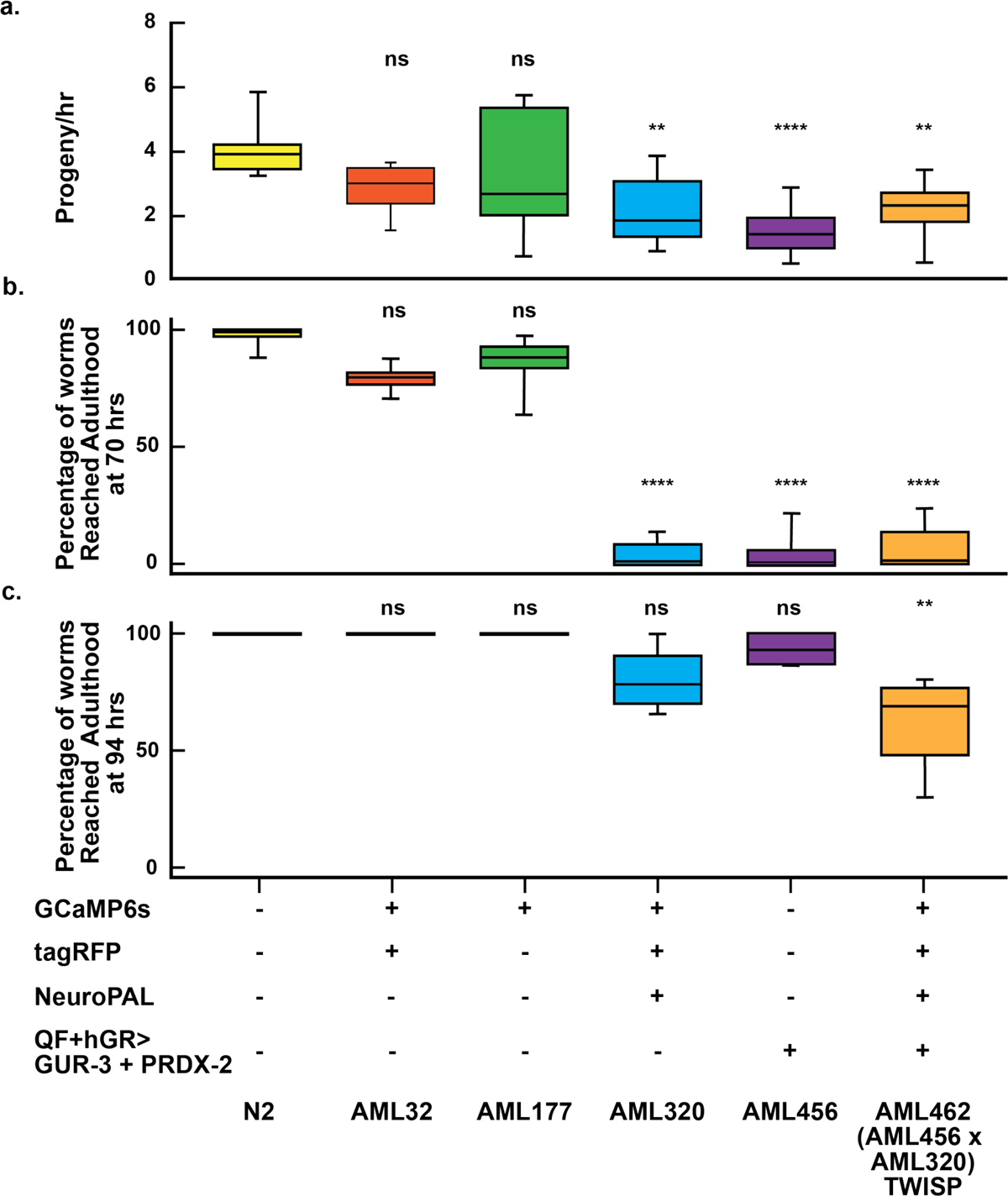
Rate of growth and progeny production decrease with transgenic load and actuator expression. a) Rate of progeny and b) percentage of animals that reach adulthood in 70 hrs or c) in 94 hrs is reported for several strains carrying various components of the TWISP system. Box shows median (line) and 25% percentile and 75% percentile (bottom and top) values, whiskers show min and max values. Exact values and number of plates are reported in Supplementary Tables S3, S4 and S5. Statistical significance is with respect to WT, using Kruskal-Wallis followed by Dunn’s multiple comparisons, *<0.03, **<0.002, ***<0.0002 & ****<0.0001.

**Figure 6.**
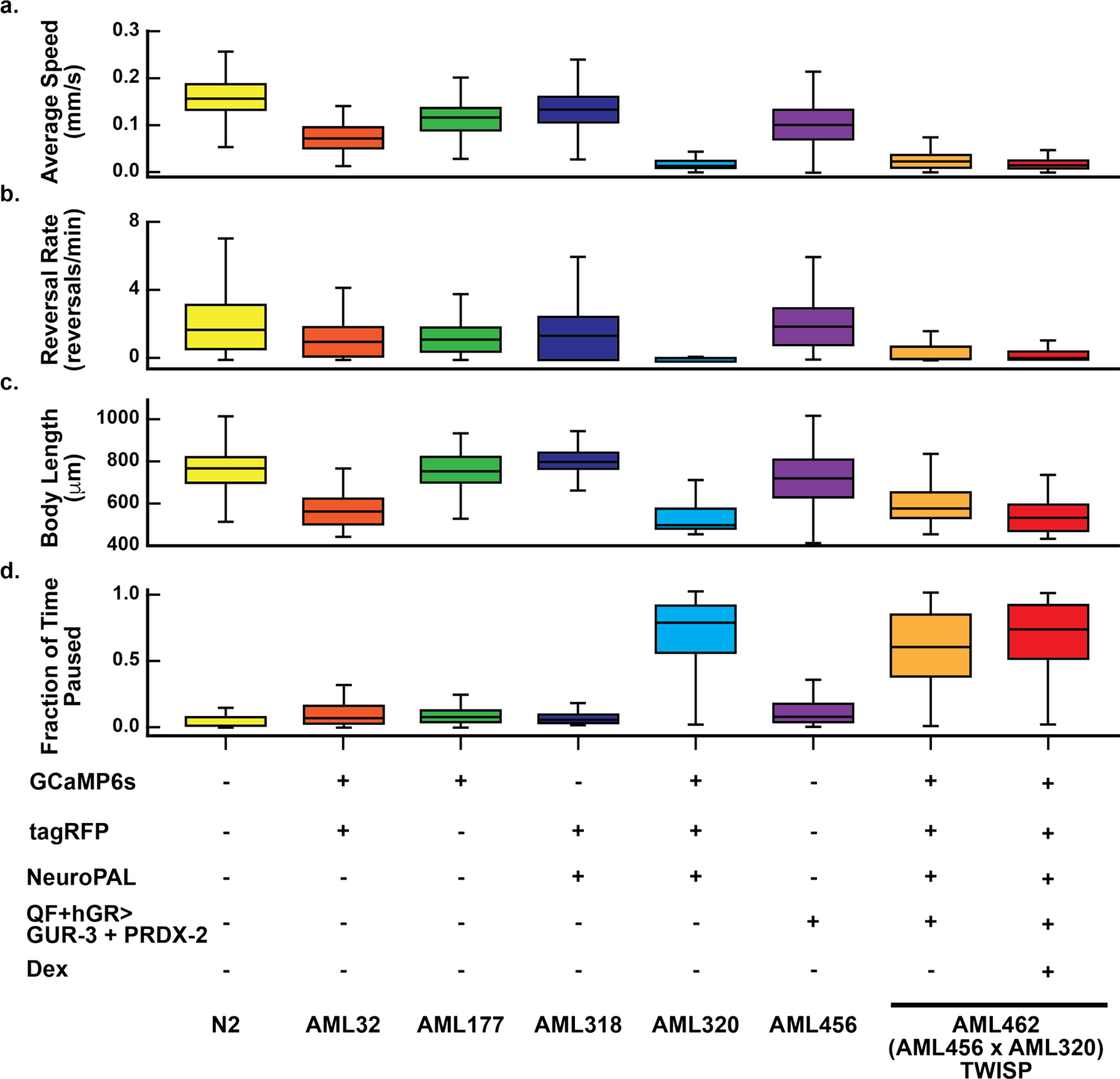
Locomotion decreases with transgenic load and actuator expression. a) Speed b) reversal rate c) body length and d) fraction of time paused is reported for animals from strains containing various components of the TWISP system. The TWISP strain is measured with and without dexamethasone treatment. The combination of NeuroPAL and GCaMP expression decreases locomotion. Box and whisker plots report the distribution of behavior across animal tracks (not plates). The number of tracks recorded per condition, from left to right, are N= [1706, 304, 1283, 1940, 393, 1374, 723, 647]. Box indicates median and interquartile range. Whiskers indicate range excluding outliers. Mean+/-SD values and number of plates are reported in Supplementary Table S6. We performed a Kolmogorov-Smirnov statistical test for all conditions compared to WT. All tests were significant (p<10^-5^) after accounting for multiple hypothesis testing via Bonferroni correction.

### Using TWISP for measuring signal propagation

To demonstrate the utility of TWISP for measuring signal propagation among neurons with known neural identities, we stimulated a selection of neurons, one neuron at a time, once a minute, while recording neural population calcium activity (Fig 7). Stimulations were performed using spatially restricted two-photon 850 nm illumination (Randi *et al*. 2022). We then identified many of the neurons that we stimulated or recorded from using the NeuroPAL color-code system. For example, we stimulated neuron AVEL and observed corresponding calcium transients in AVEL, AVJR, AWBL, AWC, RMDVL and SAAVR. This provides functional evidence to suggest that AVJR, AWBL, AWC, RMDVL and SAAVR are activated by signals originating in AVEL. TWISP therefore enables systematic perturbation mapping to investigate how signals propagate through the worm nervous system at brain scale and cellular resolution.

**Figure 7.**
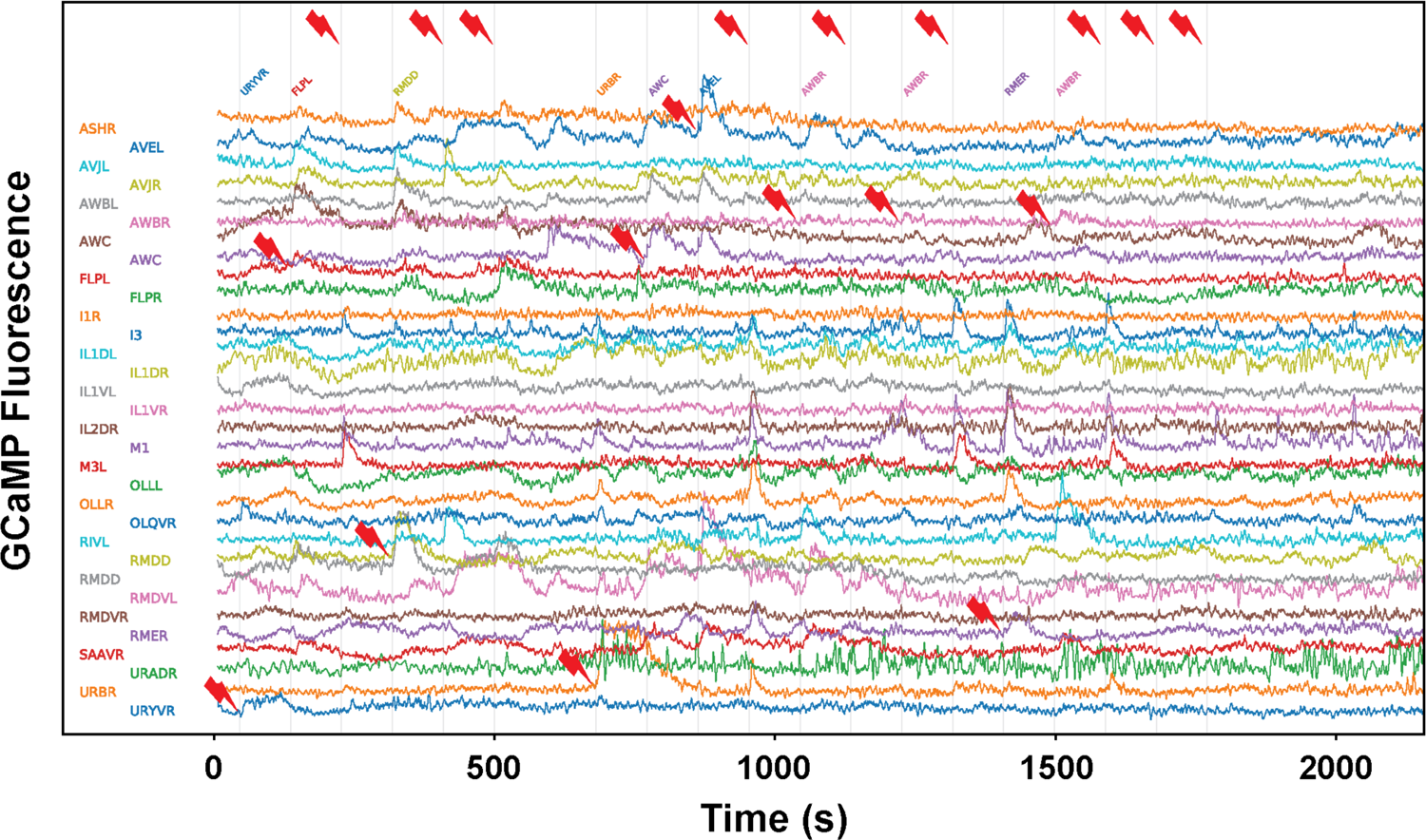
Population calcium activity in response to neural activation measured with TWISP. Calcium activity of simultaneously recorded identified neurons are shown during targeted optogenetic stimulation of individual neurons. Neuron identities are listed on the left. Gray vertical line and red thunderbolt indicate the timing and targeting of two-photon optogenetic stimulation. The name of the neuron stimulated is listed above. A thunder bolt is shown on top for those instances in which a neuron was stimulated but that neuron’s identity was not determined unambiguously. Unidentified neurons are excluded from the plot.

## DISCUSSION

In this work we presented TWISP a transgenic strain for measuring and manipulating neural activity across the brain. The key achievements of this system are 1) spectral separation of actuator and indicator that allows for calcium imaging without unwanted optogenetic activation 2) stable and robust pan-neuronal expression without lethality and 3) compatibility with a neural identification system, in this case NeuroPAL (Yemini *et al*. 2021). To our knowledge, no previous single system for all-optical neurophysiology has achieved all three of these requirements. TWISP allows for large scale measurements of how signals propagate through the brain in response to perturbations. In related work, we have used TWISP to measure a comprehensive signal propagation atlas of *C. elegans* (Randi *et al*. 2022), an endeavor that would not be possible without a system like TWISP.

TWISP makes tradeoffs to achieve its robust pan-neuronal expression with minimal spectral overlap. First, to achieve spectral separation it uses an unconventional optogenetic actuator, the GUR-3+PRDX-2 system (BHATLA AND HORVITZ 2015), that is less well characterized compared to traditionally used optogenetic proteins, such as Channelrhodopsin. GUR-3 is a *C. elegans* gustatory receptor homolog that sits in the membrane and is thought to respond to reactive oxygen species generated by light via the protein PRDX-2. PRDX-2 is thought to detect reactive oxygen species intracellularly and then activate GUR-3 at the membrane, which in turn is proposed to evoke neural activity via second messengers (Bhatla And Horvitz 2015; Quintin *et al*. 2022). The exact mechanism is unknown, however, and GUR-3 may instead act on the membrane potential directly, as a light-gated ion channel (Sonya *et al*. 2023). Regardless of the mechanism, GUR-3+PRDX-2 has already been shown to evoke light-dependent calcium transients (Bhatla And Horvitz 2015), to release glutamate to downstream synaptic partners (BHATLA *et al*. 2015), and to evoke behavioral responses (Bhatla And Horvitz 2015; Randi *et al*. 2022), all consistent with its role as an effective light-sensitive neural actuator. Notably, light-evoked GUR-3 dependent calcium transients have similar GCaMP6s fluorescent rise and fall times to those evoked via natural odor stimuli (compare, for example (Bhatla And Horvitz 2015) to (Lin *et al*. 2023). A potential advantage of GUR-3+PRDX-2 compared to a traditional opsin, is that GUR-3+PRDX-2 is activatable by light delivered intracellularly, and therefore avoids the need to illuminate the cell membrane. This is significant because it is technically more straightforward to restrict two-photon illumination to a single neuron while avoiding its neighbors when illuminating intracellularly compared to on the membrane. Intracellular illumination also avoids the need for shaping a two-photon excitation spot to match the extended shape of a cell’s membrane, which can be challenging especially in larger and irregularly shaped neurons common in mammalian systems.

TWISP also trades off the health and vitality of the animal in order to express an actuator, calcium sensor and neural identification marker in every neuron. The use of a drug-inducible system enables TWISP to be viably maintained and propagated (Fig. 4). But the resulting strain, with neural identity markers, is notably less healthy, as shown in (Fig. 5 and 6), and as expected for a strain that expresses ∼50 transgenes. We note that much of the effect on health appears to come from the combined addition of the fluorescent markers used in the NeuroPAL strain and GCaMP6s (Fig. 6). NeuroPAL animals have been reported to be less active (Yemini *et al*. 2021). Despite its non-wild-type behavior, the NeuroPAL strain is rapidly becoming a de facto standard in the field because of the value it provides in identifying neurons (Yemini *et al*. 2021; Atanas *et al*. 2022).

Evidence presented here suggests that the most severe toxicity from pan-neuronal expression of an actuator occurs early in development and can be avoided by only inducing expression in adulthood. Future investigations are needed to better understand the exact mechanism of this toxicity. And similarly, more work is needed to understand the mechanism by which fluorophores for neural identification combined with GCaMP also produce non-wild-type behavior.

TWISP should enable several new investigations that were previously inaccessible to the *C. elegans* systems neuroscience community. For example, in (Randi *et al*. 2022)we are using TWISP to compare signal propagation in wild-type and *unc-31* mutants in order to explore the role of extrasynaptic signaling in the nervous system at brain scale and cellular resolution and to validate predictions based on recent gene expression (Taylor *et al*. 2021) and peptide-receptor interaction screens (Beets *et al*. 2022; Ripoll-Sánchez *et al*. 2022). Similarly, by leveraging the powerful genetics and mutant libraries of the worm, TWISP should enable investigations into the role of specific transmitters and neuropeptides (Chase And Koelle 2007) and genes (Hobert 2013) to provide a better understanding of the molecular genetic underpinnings of the nervous system.

## Acknowledgements

F. R. contributed to this article as an employee of Princeton University and the views expressed do not necessarily represent the views of Regeneron Pharmaceuticals Inc.. Some strains were provided by the CGC, which is funded by NIH Office of Research Infrastructure Programs (P40 OD010440).

## Resources and reagents

Strains used in this work are being made available through the Caenorhabditis Genetics Center; plasmids will be made available through Addgene. Supplementary Table S1 describes strains used in this study. Supplementary Table 2 contains lists of plasmids generated for this work.

## Author contributions

A.K.S., F.R. and A.M.L. conceived the experiments, A.K.S., F.R., S.K and S.D. conducted the experiments, A.K.S., F.R., S.K and S.D analyzed the data. A.K.S and A.M.L. wrote the manuscript.

## MATERIAL AND METHODS

### Molecular cloning and plasmids

Plasmids generated in this study are listed in (Supplementary Table S2). We used a seamless cloning strategy (In-Fusion, Takara Bio USA, Inc.). Primers were synthesized from the company IDT. Clones were confirmed by sanger sequencing (AZENTA Life Sciences, USA). In brief, we first PCR amplified both back bones and inserts using PrimeSTAR GXL DNA Polymerase, a hot start, high-fidelity polymerase (Cat#R050A, Takara Bio USA, Inc.) as per manufacturer’s instructions. Both back bones and inserts were then agarose-gel purified using NucleoSpin® Gel and PCR Clean-Up columns (Takara Bio USA, Inc.). Purified fragments then mixed at prescribed molar ratio together with In-Fusion HD Cloning Kit, (Cat#639649, Takara Bio USA, Inc.) and incubated at 55 °C for 15 minutes. 2 ml of each reaction then transformed in to 100 μl Stellar competent cells (provided with In-Fusion kit) as per manufacturer’s instructions and plated on LB plates containing appropriate selectable antibiotics.

### Worm maintenance

*C. elegans* strains were maintained according using procedures described in (BRENNER 1974) with slight modifications. All worms were handled and maintained in near dark condition, at 20 °C. Worms were exposed to dim brightfield light during transferring.

### Transgenic strains

We used micro-injections for generating transient transgenic lines followed by UV integration and miniSOG mediated rapid integration methods to create integrated transgenic animals as needed (Evans 2006; Noma And Jin 2018). Detailed information regarding plasmid concentrations injected for each transgenic worm is provided in (Supplementary Table S1). Worms created for this study will be made available from Caenorhabditis Genetics Center (CGC), University of Minnesota.

### All trans retinal (ATR) and dexamethasone (dex) treatment

Worms that expressed optogenetic actuators from the rhodopsin family, such as eTsChR, were cultivated on plates containing the necessary co-factor all-trans retinal (ATR). To prepare ATR plates, we seed an NGM plate with 250 ml *E. coli* OP50 culture mixed with 1.25 μl of 100 mM ATR from stock, a day prior to treatment start time. 100 mM ATR stocks are prepared by dissolving 100 mg of ATR (Cat# R2500, Sigma-Aldrich) in 3.52ml Ethyl alcohol and then filter-sterilization using 0.2 μm filters. 100 mM ATR stocks are aliquoting and stored in smaller volume at −20 °C, in dark tubes.

Animals that express actuators under the control of the drug-inducible QF+hGR system were treated with dexamethasone (dex) to induce gene expression prior to experiments. To prepare Dex-NGM plates, we added 2 ml of dex stock solution (100 mM dex in DMSO) to each liter of NGM-agar media, 5 minutes before pouring the plates, while stirring. Dex-plates were stored at 4 °C for up to a month. Dex stocks were prepared by dissolving 1 g Dexamethasone (Cat# D1756, Sigma-Aldrich) in 25.5 ml DMSO (Cat#D8418, Sigma-Aldrich). Dex stocks were filter sterilized, aliquoted and stored at −80 °C in the dark until needed.

### Light-evoked behavior response assay

To test the behavioral response of transgenic worms expressing various pan-neuronal actuators, we scored the animal’s behavior in response to illumination in a fluorescent stereoscope (Leica-M205FA, Leica Microsystems). Young adult animals were illuminated with either 475 nm or 505 nm light. For 475 nm light, the microscope’s built in fluorescent excitation source was used (Supplementary Fig. S1 shows spectra measured via a portable spectrometer). For 505 nm light a custom built external LED (M505L4, Thorlabs) was used. Light intensities were adjusted such that the worm was illuminated with ∼1.5 mW/mm^2^ of light as measured by a power meter placed at the focal plane. Worms were illuminated for 10-20 secs and their behavior response was scored manually using the four-point scoring criteria described in the main text.

### Multi-generation growth assay

For the multi generation growth assay described in (Fig. 4) observations were recorded about the animals health and stage over a series of days. In brief, two L4 worms were transferred to either regular- or Dex-NGM plates seeded with *E. coli* OP50, three plates each trial. After three days, plates were evaluated for the presence of the most advanced stage achieved by any progeny on the plate. If needed (in case of dex-treatment), plates were further evaluated on day 4 and day 5 for the presence of L4 animals among progeny. We further transferred two-L4 worms from dex-treatment, to new regular (to recover)- and dex (to continue treatment)-NGM plates in the second round and evaluated the plates for progeny growth after three days. The stage of the animal that reached the latest developmental staged is reported for the animals in (Fig. 4). So for example, if most animals reached L3 but one animal reached L4, L4 is reported.

### Growth and progeny production assay

For measuring growth and progeny production rate, in (Fig. 5), we recorded the proportion of animals that reached adulthood in a given time starting from the egg stage, and calculated the progeny produced using a semi-synchronization method as adapted from (Kim *et al*. 2018). Briefly, three adult hermaphrodites were placed on an NGM plate seeded with *E. coli* OP50 and allowed to lay eggs for 3 hrs. Adult worms were then removed, and the eggs were allowed to grow in dark, at 20 °C to obtain age-synchronized animals. The total number of progeny and percentage of worms reached adulthood were counted at 70 hrs (when all WT N2 progeny typically reach the adult stage) and at 94 hrs (when roughly half of AML462’s progeny reached adulthood). Progeny production rate per animal was calculated by dividing the number of progeny on the plate at the end of the assay by the original number of adults that started on the plate and dividing by the number of hours elapsed. The number of plates used is reported in Supplementary Table S3, S4 and S5.

### Locomotion measurements

A high-throughput automated behavior imaging system was used to quantify attributes of the animal’s locomotion (Liu *et al*. 2018; Liu *et al*. 2022; Kumar *et al*. 2023). Age synchronized animals were obtained by bleaching gravid animals. Eggs were then left on shaker at 450 rpm overnight. The next day, the L1 larvae were placed on *E. coli* OP50 seeded NGM plates and stored in the dark at 20 °C. Once the worms reached day 1 young adult stage, behavioral assays were performed as described in (Liu *et al*. 2022). Dex treated worms were first transferred to 200 μM dex-containing NGM plates ∼20 hrs prior to measurements.

Locomotion for each strain was measured on at least 4 plates with typically 30 to 40 animals per plate. Unlike other assays which report a single metric per plate, and then average across plates, here we calculate a single behavior metric for each worm track and then average across tracks. Each track corresponds to a single worm, although one worm typically will have multiple tracks because tracks stop and start when a worm wanders out of the field of view and returns, collides with another worm, as described in.

### Signal propagation and calcium imaging measurements

Signal propagation measurements and analysis were performed as described in (Randi *et al*. 2022). Whole-brain calcium imaging was performed on immobilized TWISP animals while individual neurons were stimulated, one-at-a-time, every 60 secs, for 0.5 secs via 850 nm 2-photon laser pulses at 500 kHz repetition rate. Before calcium imaging experiments began, multi-color imaging was used to record the color of each neuron for identification via the NeuroPAL system (Yemini *et al*. 2021). After experiments were completed, neurons were segmented based on their tagRFP-T fluorescence and fluorescent calcium traces were extracted, both via an automated analysis pipeline. Neurons identities were assigned manually based on each neuron’s position and color code from NeuroPAL. Only traces for neurons that were confidentially assigned a neuron identity were included. Animals were immobilized on 10% agarose pads in 6 μl M9 buffer, with 2 μl of 100 nm polystyrene beads (Cat# 00876-15, Polysciences) and 2 μl of levamisole (from 500 μM stock, Cat# 155228, MP Biomedicals).

### Statistical Analysis

Graphs are plotted either using Prism-v.10 (GraphPad Software LLC) (Fig. 2a, Fig. 3a. Fig. 5) or Matlab (MathWorks) (Fig. 6). Kruskal-Wallis (One-way ANOVA) with Dunn’s multiple comparisons test was used for calculating statistical significance in (Fig. 5). For comparing locomotion from our high throughput behavior assay, a Kolmogorov-Smirnov test was used, and the Bonferroni multiple hypothesis correction was taken into account.

Supplementary Data: Sharma, et al., 2023.

---

**Supplementary Figure S1:**
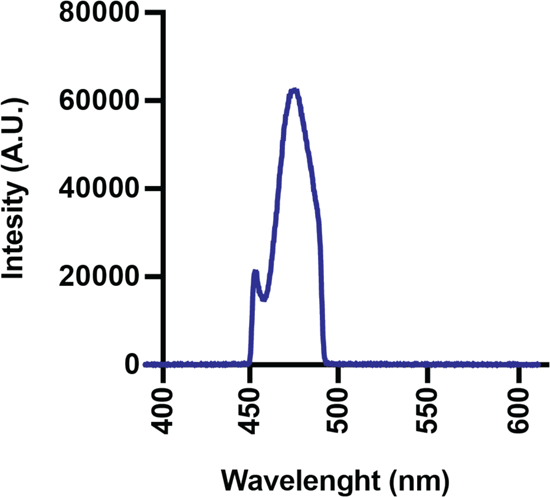
Spectra of blue light used in behavior response assays performed in Fig. 1c, Fig. 2b and Fig. 3b.

**Supplementary Table S1:**
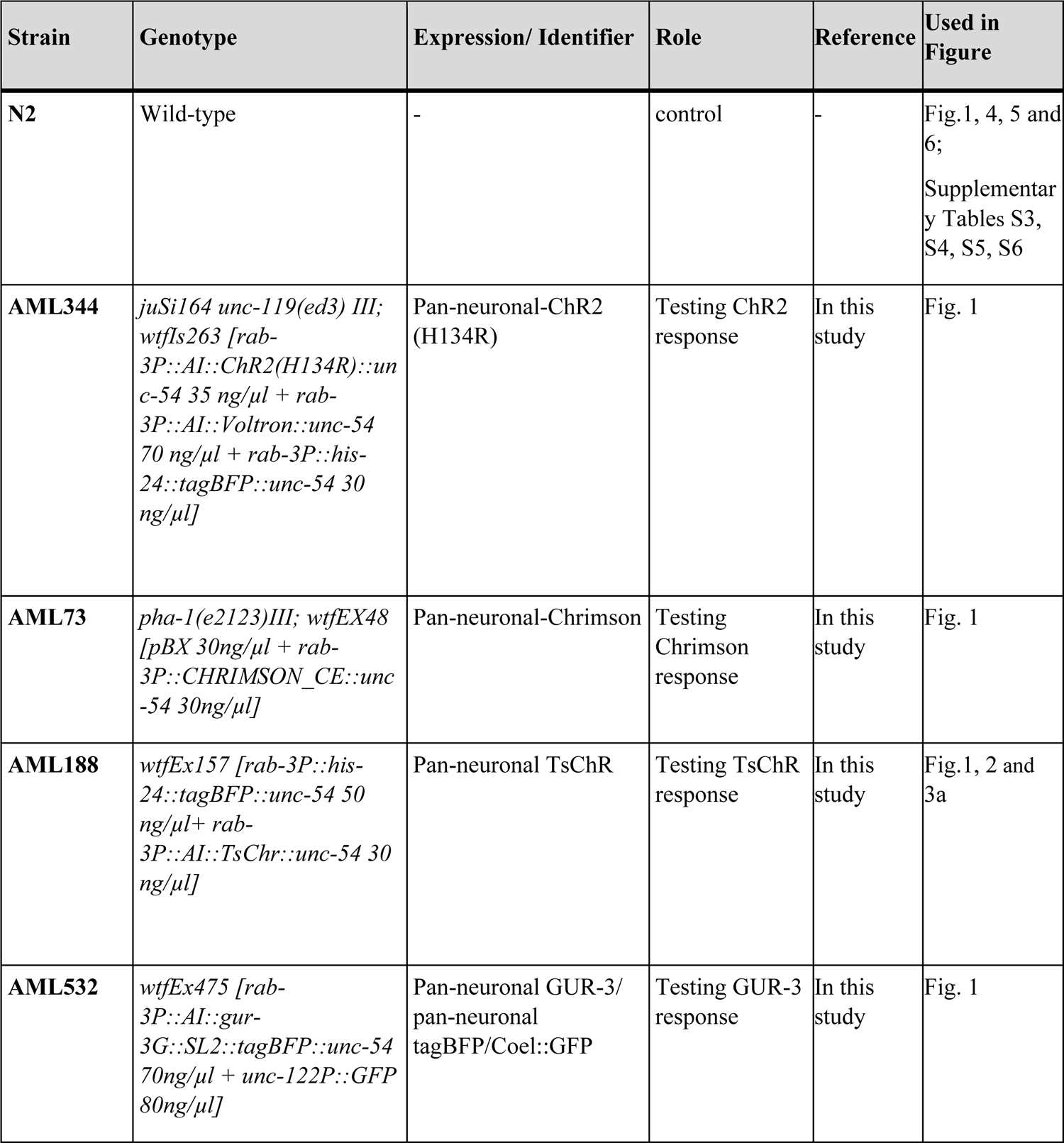
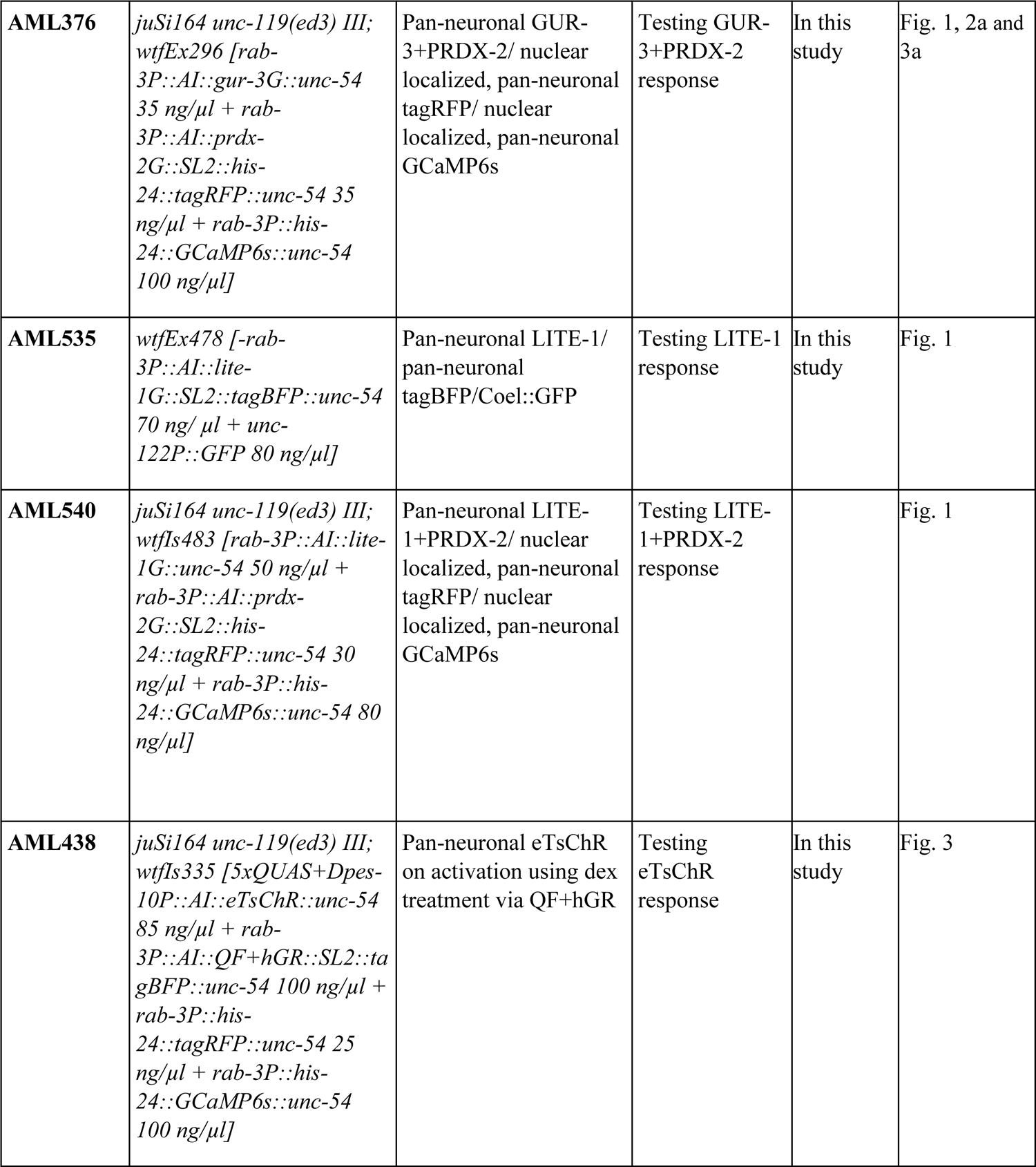
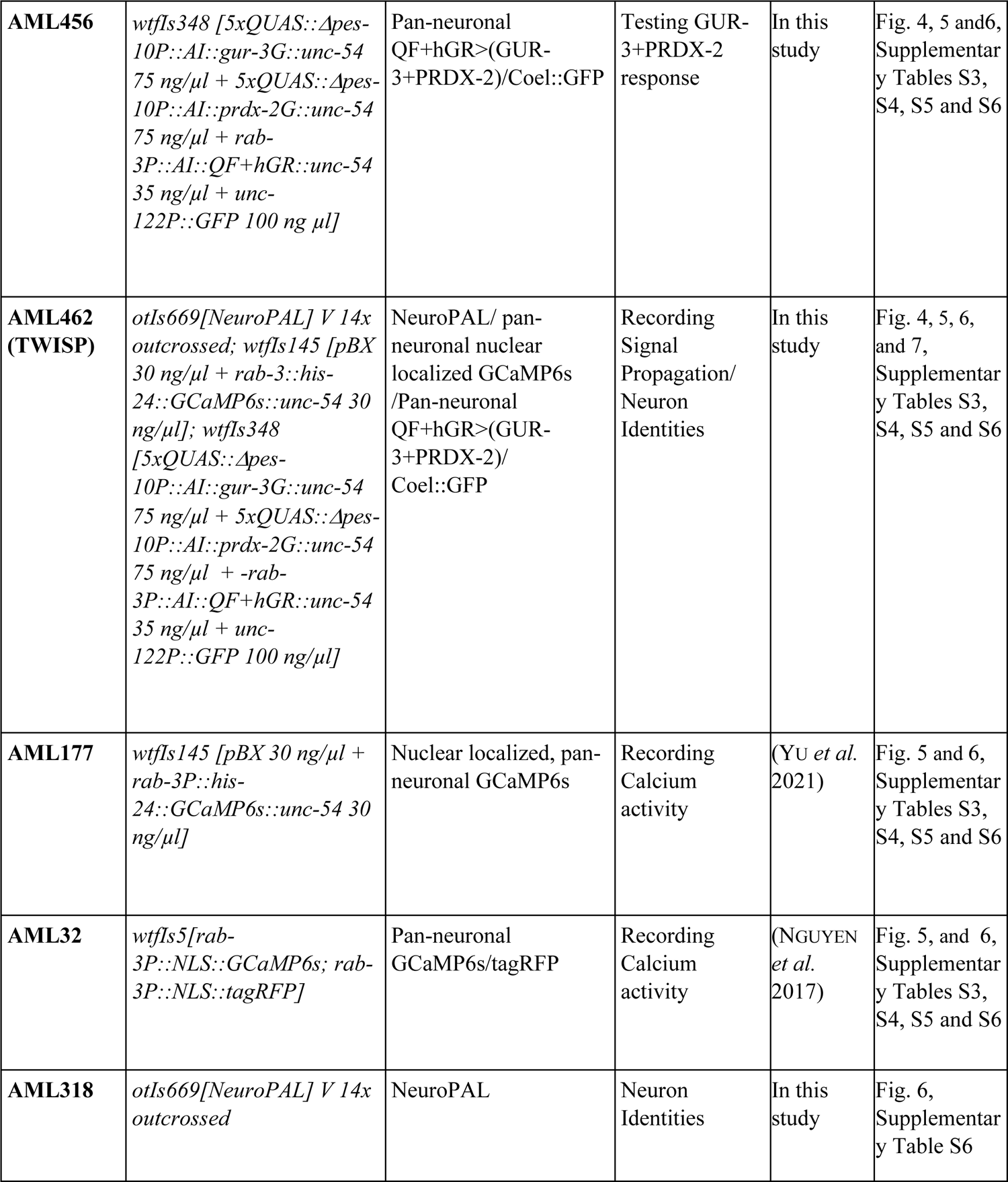
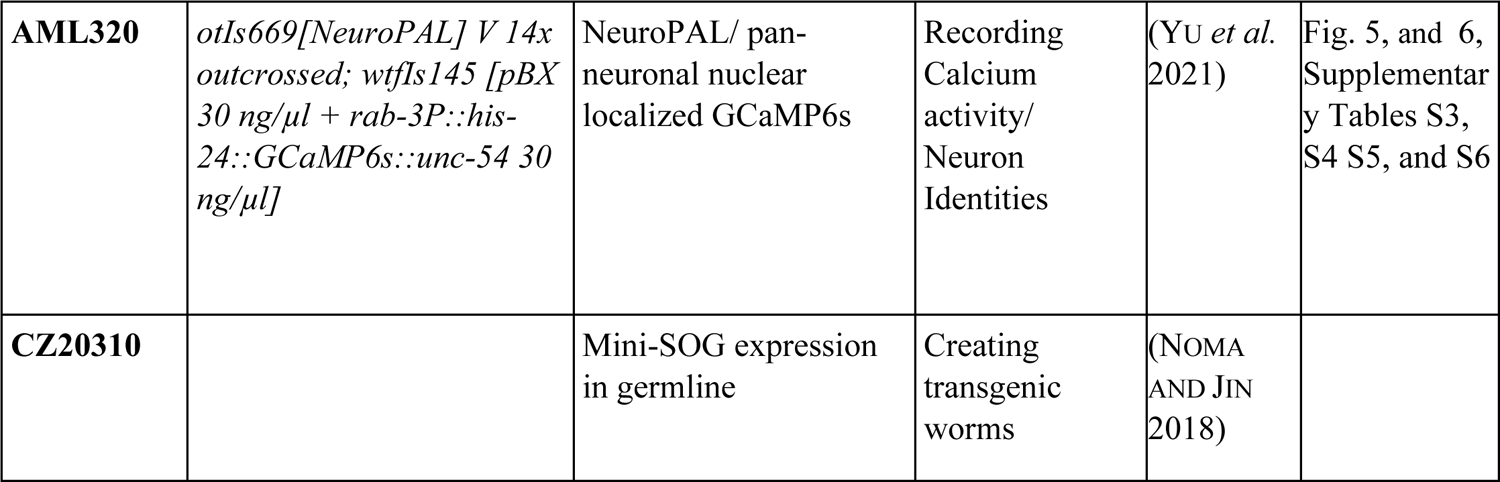
List of worms used, and transgenic worms created for this work, sorted by figure in which they first appear.

**Supplementary Table S2:** List of plasmids created for this study.

| Plasmid Created for this study | Description |
| --- | --- |
| *pAS1-rab-3P::his-24::tagRFP* | Pan-neuronal expression of nuclear localized fluorophore “tagRFP”, regulated by *rab-3* promoter and *unc-54* 3' utr |
| *pAS1-rab-3P::his-24::tagBFP* | Pan-neuronal expression of nuclear localized fluorophore “tagBFP”, regulated by *rab-3* promoter and *unc-54* 3' utr |
| *pAS1-rab-3P::his-24::GCaMP6s* | Pan-neuronal expression of nuclear localized calcium sensor “GCaMP6s”, regulated by *rab-3* promoter and *unc-54* 3' utr |
| *pAS3-rab-3P::AI::ChR2(H134R)* | Pan-neuronal expression of opsin “ChR2(H134R)”, regulated by *rab-3* promoter and *unc-54* 3' utr |
| *pAS3-rab-3P::AI::Voltron* | Pan-neuronal expression of voltage sensor “Voltron”, regulated by *rab-3* promoter and *unc-54* 3' utr |
| *pAS3-rab-3P::CHRIMSON_CE* | Pan-neuronal expression of opsin “Chrimson”, regulated by *rab-3* promoter and *unc-54* 3' utr |
| *pAS3-rab-3P::AI::TsChr* | Pan-neuronal expression of opsin “TsChR”, regulated by *rab-3* promoter and *unc-54* 3' utr |
| *pAS3-rab-3P::AI::gur-3G* | Pan-neuronal expression of “GUR-3” using a genomic fragment, regulated by *rab-3* promoter and *unc-54* 3' utr |
| *pAS3-rab-3P::AI::lite-1G* | Pan-neuronal expression of “LITE-1” using a genomic fragment, regulated by *rab-3* promoter and *unc-54* 3' utr |
| *pAS3-rab-3P::AI::QF+hGR::SL2::tagBFP* | Pan-neuronal expression of chimeric protein “QF+hGR” and fluorophore “tagBFP”, regulated by *rab-3* promoter and *unc-54* 3' utr |
| *pAS3-rab-3P::AI::gur-3G::SL2::tagBFP* | Pan-neuronal expression of “GUR-3” using a genomic fragment and fluorophore “tagBFP”, regulated by *rab-3* promoter and *unc-54* 3' utr |
| *pAS3-rab-3P::AI::prdx-2G::SL2::his-24::tagRFP* | Pan-neuronal expression of “PRDX-2” using a genomic fragment and fluorophore “tagRFP”, regulated by *rab-3* promoter and *unc-54* 3' utr |
| *pAS3-rab-3P::AI::lite-1G::SL2::tagBFP::unc-54* | Pan-neuronal expression of “LITE-1” using a genomic fragment and fluorophore “tagBFP”, regulated by *rab-3* promoter and *unc-54* 3' utr |
| *pAS3-rab-3P::AI::QF+hGR* | Pan-neuronal expression of chimeric protein “QF+hGR”, regulated by *rab-3* promoter and *unc-54* 3' utr |
| *pAS3-5xQUAS+Δpes-10P::AI::eTsChR* | Expression of opsin “eTsChR” on activation via “QF+hGR” chimeric protein |
| *pAS3-5xQUAS::Δpes-10P::AI::gur-3G* | Expression of “GUR-3” on activation via “QF+hGR” chimeric protein |
| *pAS3-5xQUAS::Δpes-10P::AI::prdx-2G* | Expression of “PRDX-2” on activation via “QF+hGR” chimeric protein |

**Note:** pAS1 and pAS3 plasmid backbones are including *unc-54* 3' utr sequence.

**Supplementary Table S3:**
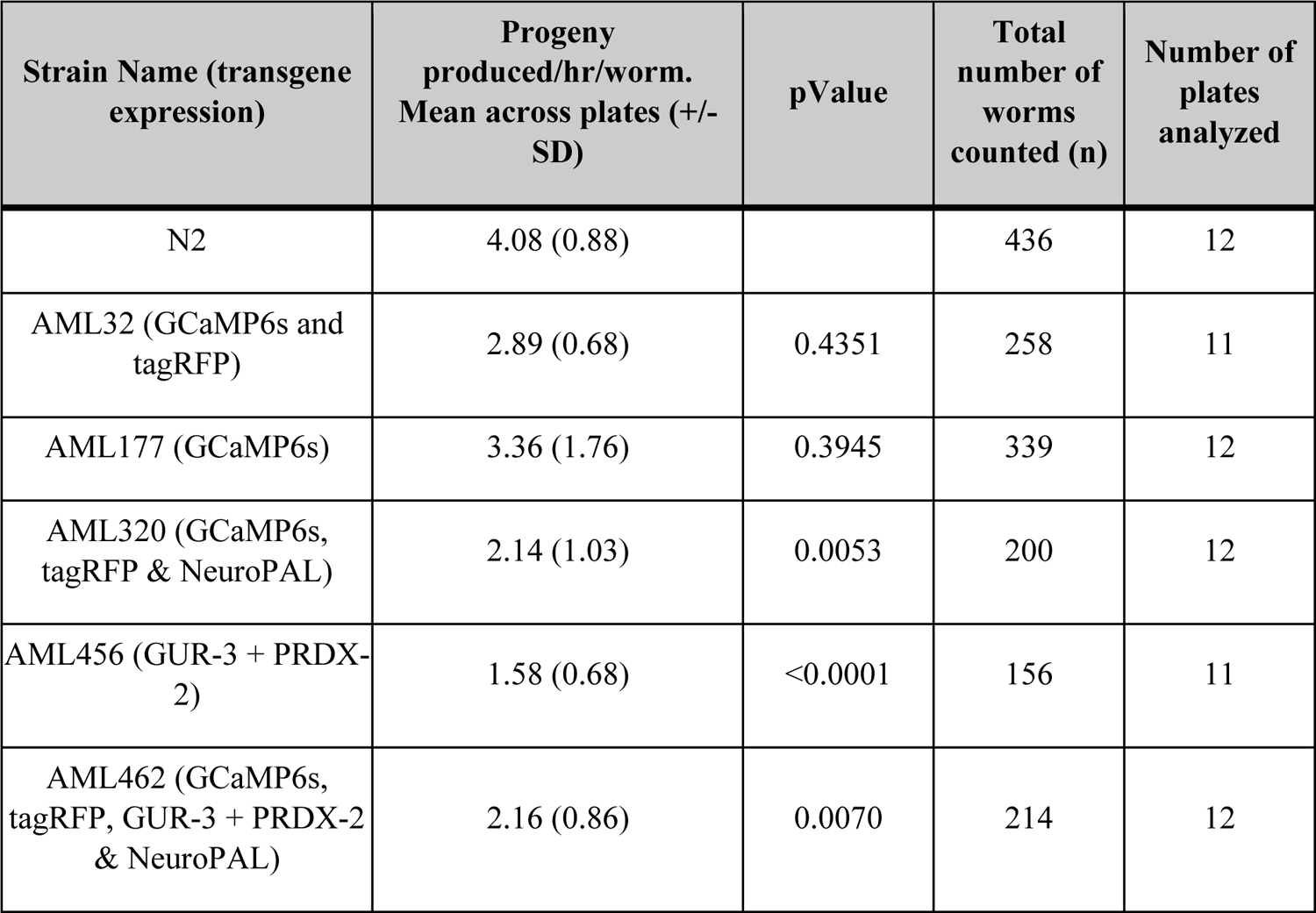
Data from progeny production assay, corresponding to Figure 5a, including number of plates, total number of worms counted and statistical values.

**Supplementary Table S4:** Data from growth assay at 70 hrs, corresponding to Figure 5b, including number of plates, total number of worms counted and statistical values.

| Strain Name (transgene expression) | Percentage of worms reached to Adulthood at ~70 hrs: Mean across plates (+/- SD) | pValue | Total number of worms counted (n) | Number of plates analyzed | Remarks |
| --- | --- | --- | --- | --- | --- |
| N2 | 98.2 (3.633) |  | 436 | 12 | All adults and very few eggs |
| AML32 (GCaMP6s and tagRFP) | 80.05 (4.985) | 0.0744 | 258 | 11 | Adults and L4s |
| AML177 (GCaMP6s) | 86.56 (9.155) | 0.6369 | 339 | 12 | Adults and L4s |
| AML320 (GCaMP6s, tagRFP & NeuroPAL) | 3.977 (5.528) | <0.0001 | 200 | 12 | Populations varies from L2s-L4s and few yAds |
| AML456 (GUR-3 + PRDX-2) | 3.254 (6.864) | <0.0001 | 156 | 11 | Most worms grew up to L3, few L4s and few yAds |
| AML462 (GCaMP6s, tagRFP, GUR-3 + PRDX-2 & NeuroPAL) | 5.737 (8.414) | <0.0001 | 214 | 12 | Populations varies from L2s-L4s and few yAds |

**Supplementary Table S5:**
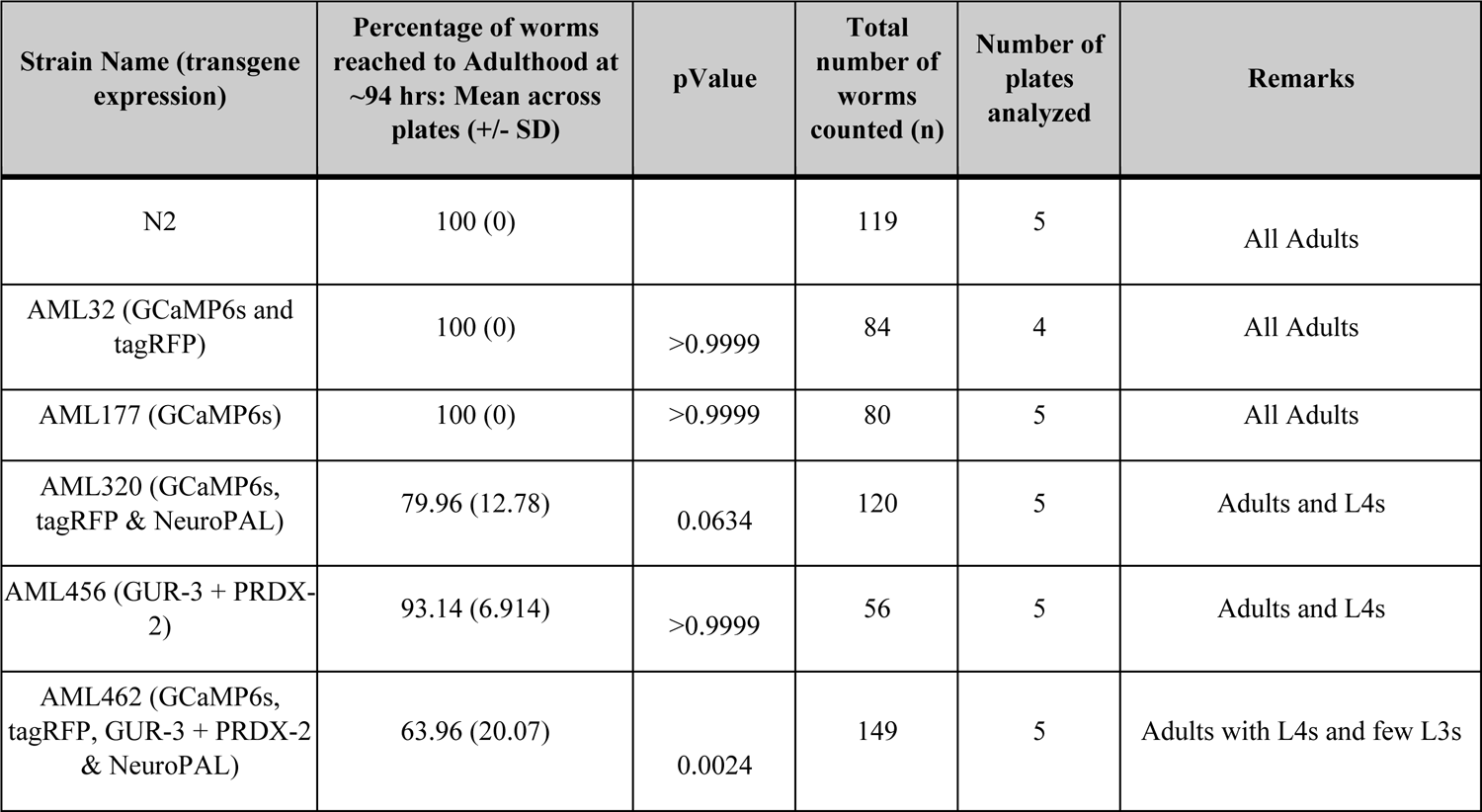
Data from growth assay at ∼94 hrs, corresponding to Figure 5c, including number of plates, total number of worms counted and statistical values.

**Supplementary Table S6:** Data from behavior analysis, corresponding to Figure 6, including number of plates recorded, total number of tracks in analysis and statistical values for each strain for each parameter.

| Strain Name (transgene expression) | Average Speed (mm/sec) Mean (+/- SD), pValue | Reversal Rate, (reversals/min) Mean (+/- SD), pValue | Length (mm) Mean (+/- SD), pValue | Fraction of Time Paused. Mean (+/- SD), pValue | Number of Tracks analyzed | Number of assays (plates) |
| --- | --- | --- | --- | --- | --- | --- |
| N2 | 0.15 (0.042) | 2 (1.6) | 752.57 (110.1) | 0.064 (0.12) | 1706 | 12 |
| AML32 (GCaMP6s and tagRFP) | 0.073 (0.023), 2.27<sup>-163</sup> | 1.2 (1.2), 3.48<sup>-15</sup> | 565.14 (74.66), 9.14<sup>-128</sup> | 0.11 (0.13), 5.22<sup>-19</sup> | 304 | 4 |
| AML177 (GCaMP6s) | 0.11 (0.032), 2.75<sup>-138</sup> | 1.2 (0.96), 7.79<sup>-47</sup> | 745.98 (88.62), 2.52<sup>-5</sup> | 0.084 (0.077), 6.48<sup>-51</sup> | 1283 | 8 |
| AML318 (tagRFP & NeuroPAL) | 0.13 (0.041), 7.04<sup>-53</sup> | 1.59 (1.55), 1.45<sup>-10</sup> | 791.83 (66.86), 1.14<sup>-52</sup> | 0.061 (0.072), 7.07<sup>-6</sup> | 1940 | 4 |
| AML320 (GCaMP6s, tagRFP & NeuroPAL) | 0.017 (0.011), 6.59<sup>-265</sup> | 0.17 (0.56), 1.90<sup>-134</sup> | 536.84 (88.38), 2.72<sup>-143</sup> | 0.7 (0.24), 4.53<sup>-243</sup> | 393 | 4 |
| AML456 (GUR-3 + PRDX-2) | 0.097 (0.043), 6.08<sup>-184</sup> | 1.9 (1.6), 0.0211 | 709.32 (125.44), 4.12<sup>-26</sup> | 0.15 (0.22), 7.78<sup>-51</sup> | 1374 | 4 |
| AML462 (GCaMP6s, tagRFP, GUR-3 + PRDX-2 & NeuroPAL) | 0.026 (0.021), <1<sup>-324</sup> | 0.52 (1), 2.38<sup>-110</sup> | 595.19 (93.33), 2.05<sup>-173</sup> | 0.59 (0.29), 1.644<sup>-318</sup> | 723 | 8 |
| AML462 (on-dex) | 0.019 (0.013), <1<sup>-324</sup> | 0.46 (1), 2.83<sup>-122</sup> | 536.98 (91.75), 1.88<sup>-239</sup> | 0.69 (0.25), <1<sup>-324</sup> | 647 | 4 |

